# Soleus muscle weakness in Cerebral Palsy: muscle architecture revealed with Diffusion Tensor Imaging

**DOI:** 10.1101/436485

**Authors:** Annika S. Sahrmann, Ngaire Susan Stott, Thor F. Besier, Justin W. Fernandez, Geoffrey G. Handsfield

## Abstract

Cerebral palsy (CP) is associated with movement disorders and reduced muscle size. This latter phenomenon has been observed by computing muscle volumes from conventional MRI, with most studies reporting significantly reduced volumes in leg muscles. This indicates impaired muscle growth, but without knowing muscle fiber orientation, it is not clear whether muscle growth in CP is impaired in the along-fiber direction (indicating shortened muscles and limited range of motion) or the cross-fiber direction (indicating weak muscles and impaired strength). Using Diffusion Tensor Imaging (DTI) we can determine muscle fiber orientation and construct 3D muscle architectures to examine along-fiber length and cross-sectional area separately. Such an approach has not been undertaken in CP. Here, we use advanced DTI sequences with fast imaging times to capture fiber orientations in the soleus muscle of children with CP and age-matched, able-bodied controls. Physiological cross sectional areas (PCSA) were reduced (37 ± 11%) in children with CP compared to controls, indicating impaired muscle strength. Along-fiber muscle lengths were not different between groups, but we observed large variance in length within CP group. This study is the first to demonstrate functional strength deficits using DTI and implicates impaired cross-sectional muscle growth in children with cerebral palsy.

## INTRODUCTION

Cerebral Palsy (CP) is one of the most common movement disorders in children. It is a disabling neuromusculoskeletal condition associated with a non-progressive neurological lesion in the brain caused before or during birth or up to two years after birth(1). Individuals with CP have impaired movements and hypertonia, which can limit both physical activity and social participation(2,3). Although the neural lesion is non-progressive(4), the biomechanical impairments in CP are progressive, becoming worse as the child grows and ages(5). Since the progression of CP appears to be musculoskeletal and not neural, this disorder may be greatly illuminated with further investigations into musculoskeletal growth in CP.

Previous studies(4,6–11) have reported wide-spread volume deficits of the lower extremity muscles in children with CP when compared to age-matched typically developing (TD) controls. Among these observations, the soleus muscle seems to be especially affected by reduced muscle volume(6). The soleus is the largest muscle of the triceps surae and plays an essential role in standing and walking(12,13). Alterations in soleus function will thus have profound implications for stability in gait and ambulation. While muscle volume deficits can be determined relatively easily from medical imaging, these measurements do not indicate whether the muscle is short in the along-fiber direction or small in the cross-fiber direction. Indeed, knowledge of muscle fiber orientation is necessary to determine this. Functionally, muscle size in the cross-fiber direction can be expressed as the physiological cross-sectional area (PCSA) and is related to the muscle’s force generating capacity(14). Fascicle length is the length of the muscle in the along-fiber direction and is related to the range of motion and contraction velocity of a muscle. To calculate PCSA and fascicle length from medical imaging, specific fiber architectures must be known. Since CP often presents heterogeneously across subjects, ‘typical’ fiber architecture across subjects is not expected and determination of subject-specific fiber orientation is important. With such knowledge, muscle architecture characteristics such as pennation angle, fascicle length, and PCSA can be calculated to help provide information about functional impairments at the muscle level in children with CP. This information may inform orthopaedic surgeries such as tendon lengthenings, which specifically target a muscle-tendon unit and can be tailored or ruled out with knowledge of the length and cross-sectional area of the muscle being targeted.

One *in vivo* technique for acquiring muscle fiber data is Diffusion Tensor Imaging (DTI) which is a Magnetic Resonance Imaging (MRI) sequence that encodes diffusion to evaluate directions of fluid motion within tissues(15,16). Since water diffuses preferentially along fibers rather than across them in skeletal muscle, DTI can be used to reconstruct and analyze skeletal muscle fiber tracts(17–21), which can then be used to calculate muscle architecture. The long imaging times of DTI have prevented its use in CP populations in the past, since it is essential that patients remain still for the length of the scanning. Advancements in DTI have made it possible to acquire images quickly, opening up the possibility to use this technology in CP patients.

The purpose of this study was to investigate and compare the muscle fiber architecture of the soleus muscles in children with and without CP, focusing on the PCSA, fascicle length, and pennation angle. By beginning with a foundational understanding of the muscle architecture in CP and how it differs from TD muscle architecture, the muscular contribution to impaired biomechanics in CP can be illuminated. With muscle fiber architecture, predictions about functional impairments such as reduced strength, fatigue, and impaired gait of children with CP can be more reliable and rehabilitation therapies can be evaluated in terms of muscle architecture.

## MATERIALS AND METHODS

### Participant Characteristics and Imaging

Images were collected from 9 volunteers with CP, ranging from level I to III in the Gross Motor Function Classification System(22,23) (GMFCS) with the following characteristics:. [mean ± SD (range)]: age: 11.1 ± 2.0 (8-13) years, height: 145.1 ± 10.8 (125.0-154.0) cm, body mass: 37.7 ± 10.3 (22.0-51) kg, body mass index: 17.6 ± 2.9 (14.1-21.5) kg/m². Participant characteristics are provided in Table 1. Inclusion criteria included the ability to safely undergo MRI and remain motionless in the MRI scanner for the duration of the imaging time. Nine age- and gender-matched typically developed (TD) participants were recruited for comparison as a control group, with the following characteristics: [mean ± SD (range)]: age: 11.1 ± 2.0 (8-13) years, height: 150.6 ± 11.8 (134.0-164.0) cm, body mass: 39.6 ± 8.2 (27.0-51.0) kg, body mass index: 17.3 ± 1.5 (15.0-20.2) kg/m². The study protocol was approved by the University of Auckland Human Participants Ethics Committee. Parents/guardians of participants provided informed consent prior to study participation and all participants assented to the study.

**Table 1:** Characteristics for all CP subjects

|  | CP 1 | CP 2 | CP 3 | CP 4 | CP 5 | CP 6 | CP 7 | CP 8 | CP 9 |
| --- | --- | --- | --- | --- | --- | --- | --- | --- | --- |
| Age (years) | 13 | 11 | 12 | 13 | 12 | 8 | 9 | 8 | 8 |
| Gender (M/F) | M | F | F | M | M | M | M | F | F |
| Height (m) | 1.50 | 1.48 | 1.54 | 1.57 | 1.60 | 1.25 | 1.36 | 1.14 | 1.17 |
| Weight (kg) | 37 | 31 | 49 | 41 | 51 | 22 | 33 | 17 | 20 |
| BMI (kg/m<sup>2</sup>) | 16.4 | 14.2 | 20.7 | 16.6 | 19.9 | 14.1 | 17.8 | 13.1 | 14.6 |
| GMFCS | II | II | I | III | III | II | I | II | I |

Imaging data were collected from knee to ankle. The scanning was conducted on a 3.0T Siemens Skyra Scanner (Erlangen, Germany) using a high-resolution 3D T1 VIBE Dixon sequence with the following parameters: TE/TR: 5.22 ms/ 10.4 ms; FOV: 128mm × 228mm × 384mm [R-L, A-P, S-I]; spatial resolution: 0.8 mm × 0.8 mm × 0.8 mm; imaging time: 5min. After the Dixon scan, we immediately scanned using an echo planar imaging sequence with the following parameters: TE/TR/α: 74.0 ms/ 4400 ms; b-value: 500 mm/s^2^; FOV: 200mm × 200mm in-plane; axial slice thickness: 5 mm; in-plane spatial resolution: 1.64 mm × 1.64 mm; two stacks of 35 images; imaging time: 6min. The subjects were scanned in feet first supine position with 10° of knee flexion. The subjects’ ankles were positioned in a neutral orientation using a foam block and bean bag. An accessory flex coil and the body coil were used. DTI image stacks were merged with a custom code written in Matlab R2016a (Natick, Massachusetts, USA). Post-processing was conducted using DSI Studio(24).

### Image Processing

To increase image signal, T1 images were resampled from a slice thickness of 0.8mm to 4mm using Matlab (R2016a). We segmented three characteristic regions of the soleus muscle consistent with regions identified from anatomical studies(25,26) (Fig 1). Segmentation was conducted on T1 images in the axial plane using ITK Snap(27) (V 3.4.0). T1-weighted images were spatially registered to the DT image space and the segmented muscle regions were exported as binary masks for DTI post processing. Fiber tracking was conducted by importing the transformed binary mask into the DSI studio software, and using the fiber tracking algorithm(28) for single-thread DTI implementation with the following parameters: angular threshold: 40°, FA: 0.1, minimum tract length: 20mm, maximum tract length: 200mm. After this process, DTI tracts were visually inspected to confirm that they originated and terminated near aponeurosis locations. Muscle fascicle lengths were then taken as the DTI tract length.

**Fig 1:**
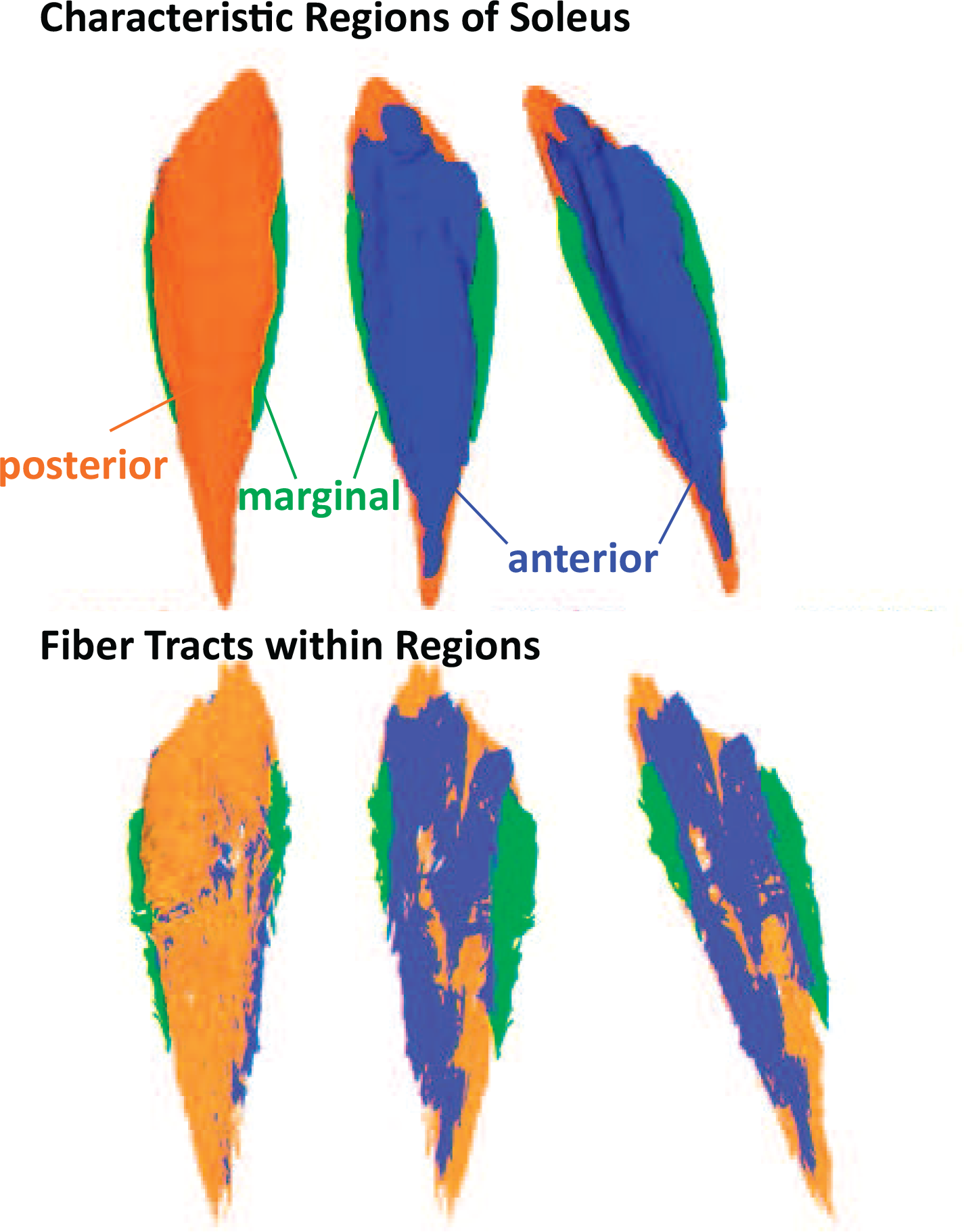
3D reconstructions of segmented MRI images illustrate the 3 characteristic regions of the soleus muscle—posterior (orange), anterior (blue) and marginal (green). Fiber tracts reconstructed from DTI within those masks represent unique fiber regions within the muscle.

### PCSA

PCSAs for each soleus were calculated from the soleus fascicle lengths using the following equation(29,30). 
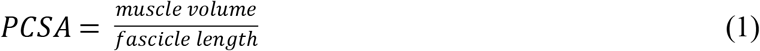
where *PCSA* is the physiological cross-sectional area, *muscle volume* is the volume of the soleus acquired from image post-processing in units of *cm*^*3*^ and fascicle length is the length of a fiber obtained from DTI post-processing software in *cm*.

### Pennation Angle

The conventional definition of pennation angle is the 2D-angle between the muscle fiber and the line of action of the muscle(31,32). The 2D nature of this definition makes it difficult to apply to complex three dimensional data. Another definition describing 3D angles has been offered in the past(25,33). Consistent with these, pennation angles were calculated as the angle between the tangent vector of the fiber bundle and the norm vector of the muscle surface at the insertion point of the bundle. Since each region of the muscle consists of multiple differently arranged fiber bundles, one characteristic fiber bundle was extracted for each region and the pennation angle was computed for this fiber bundle for the region from which it was extracted. To have a reasonable comparison of angles, bundles were inspected to ensure consistency of location across subjects.

### Normalization of Variables

To reduce the effects of body height and mass on differences in muscle size and architecture, we normalized computed parameters according to the following equations. Muscle volume and PCSA were normalized to body size based on findings from a previous study(34). 
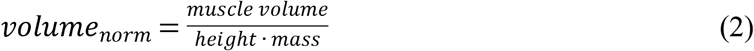
where *volume*_*norm*_ is the normalized muscle volume of the soleus, *muscle volume* is the volume of the soleus acquired from image processing in units of *cm*^*3*^, *height-mass* is the product of subject height in *cm* and body mass in *kg*. 
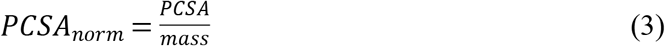
where *PCSA*_*norm*_ is the normalized physiological cross-sectional area in *cm*^*2*^*/kg*, *PCSA* is the physiological cross-sectional area of the soleus in *cm*^*2*^ and *mass* is the subject’s body mass in *kg*.

Muscle length and fascicle lengths were normalized by height as follows: 
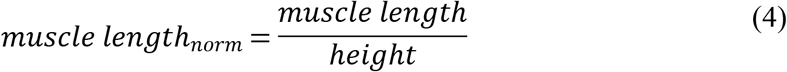

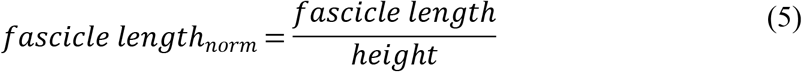
where *muscle length* is the superior-inferior length of the muscle in *cm*, *fascicle length* is the tract length obtained from DTI post-processing in *cm* and *height* is the subject’s body height in *cm*.

### Statistics

Since normalized parameters were being compared between groups, nonparametric statistical tests were necessary. For all tests of significance, the Wilcoxon ranksum test was used.

## RESULTS

### Muscle volumes and muscle lengths

Muscle volumes differed significantly between the CP and the TD group (Fig 2A). Absolute muscle volumes were 42.6% smaller in the CP group (p =0.004). Body size normalized volumes were 35.2% smaller on average for CP subjects (p =0.002). Differences in muscle length (Fig 2B) did not reach significance for either absolute or normalized lengths: absolute length difference 8.5% (p =0.131), normalized length difference 6.0% (p =0.340).

**Fig 2:**
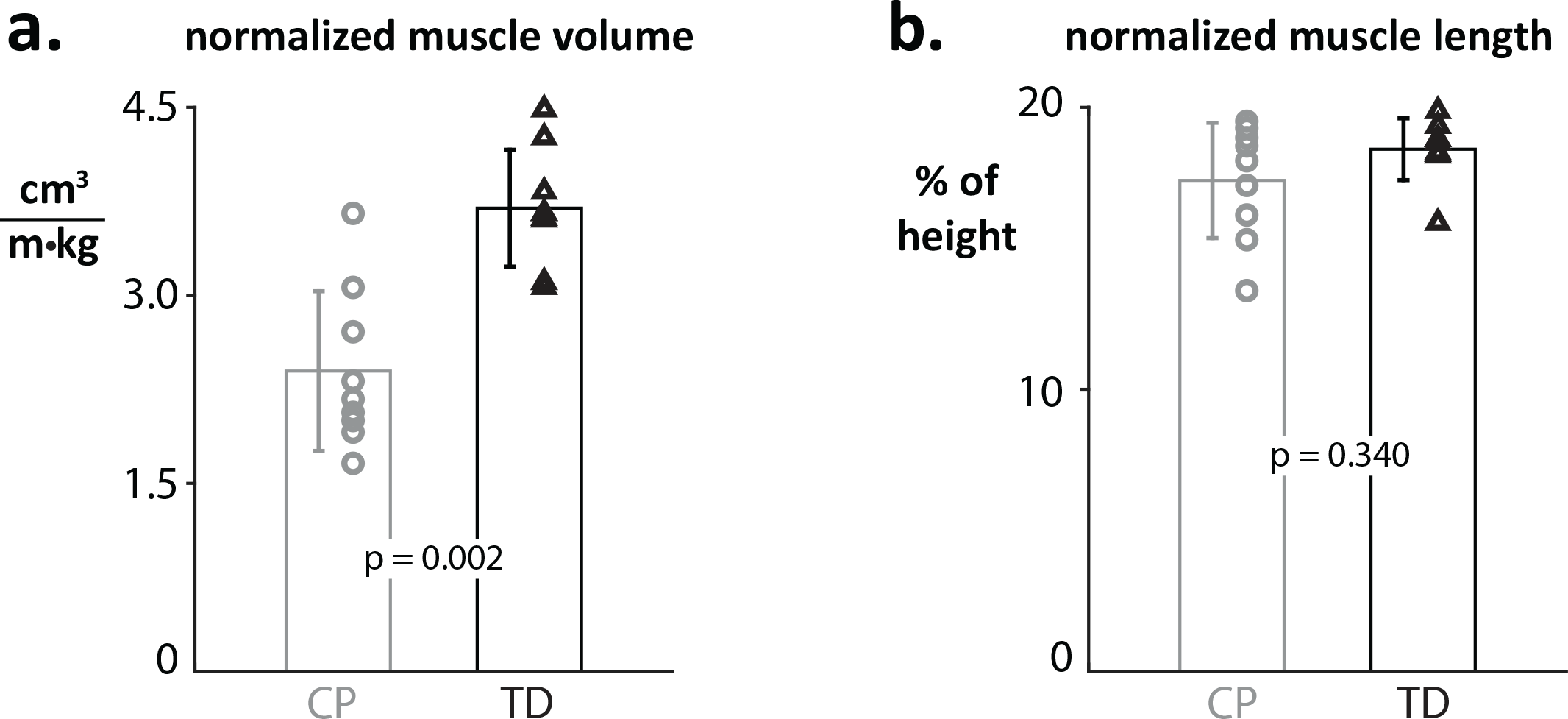
A) Normalized muscle volumes are significantly reduced in the CP cohort. B) Normalized muscle length is longer in the TD population but this difference does not reach significance.

We observed differences in the shape of the soleus muscle between the CP and TD group and within the CP group. Overall, the fiber models show that CP group has smaller muscles. Also, the subjects with CP show a less dense ‘packing’ of muscle fibers in their muscles than the subjects of the TD group, suggestive of a larger fraction of intramuscular connective tissue. CP muscles were longer and thinner compared to their TD counterparts.

The CP cohort presented smaller volumes than the TD cohort for each of the three functional regions of the soleus. The marginal region was the most reduced region and was 56.5% smaller in the CP group (p = 0.001), the anterior region was 46.1% smaller in the CP group (p = 0.004), and the posterior region was 35.5% smaller in the CP group (p = 0.019).

### Fascicle lengths and PCSA

A comparison of the median fascicle lengths between the CP and TD groups revealed slight differences in fascicle length between the cohorts (Fig 3). For the marginal and posterior compartment non-significant differences of 0.4% (p=0.465) and 2.5% (p =0.712) in the CP group were observed; fascicle lengths in the anterior compartment showed non-significant differences of 4.2% (p = 0.846) in the TD group. Absolute fascicle lengths were not significantly different in any compartment between the groups.

**Fig 3:**
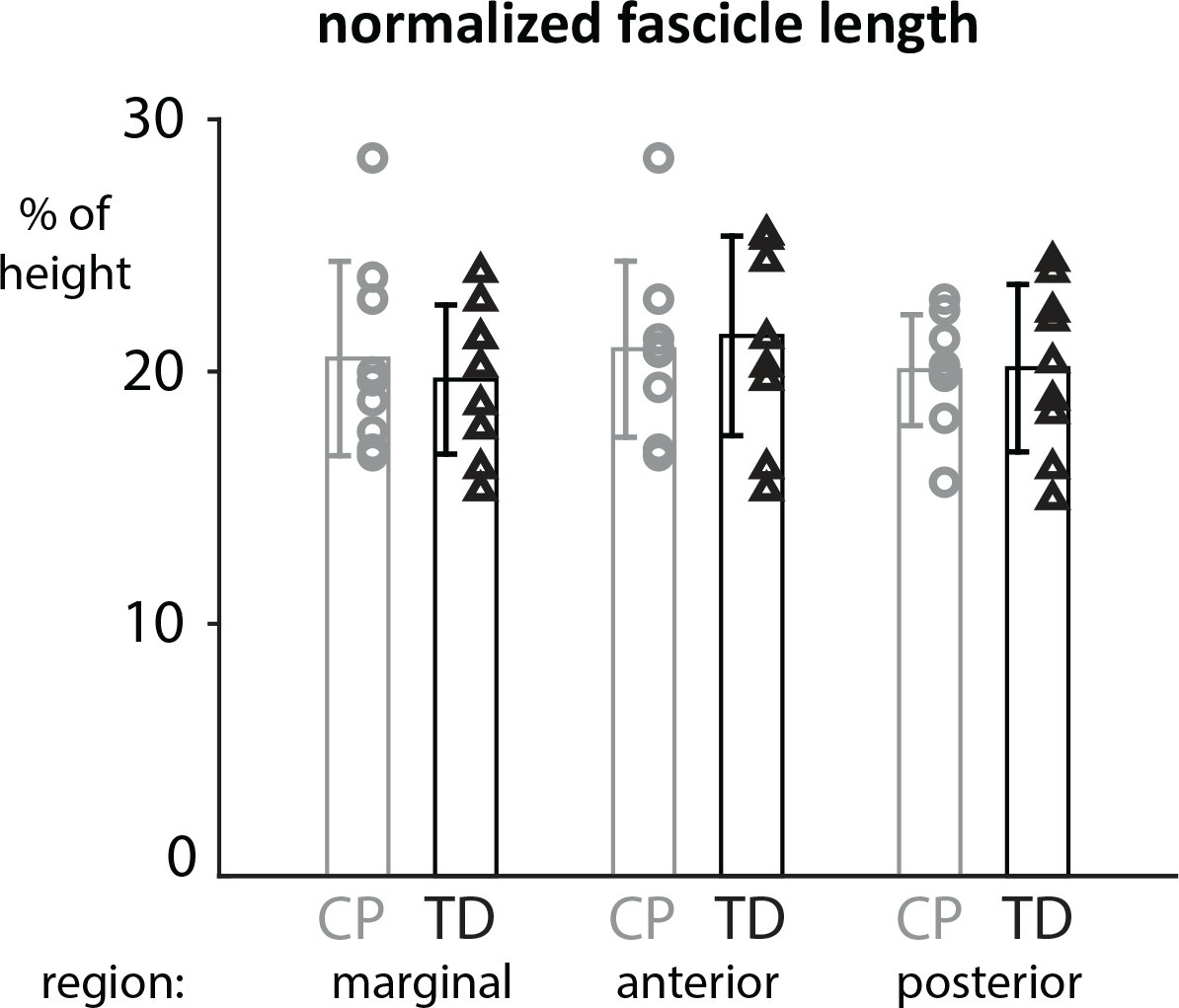
Normalized fascicle lengths in the three regions of the soleus show heterogeneities within and between groups but are not significantly different between CP and TD cohorts.

We observed heterogeneities in fascicle lengths within both cohorts (Fig 3). For instance, both the longest and the shortest normalized median fascicle length could be found within the TD group (longest 34mm with a height of 1.34m; shortest 24.5mm with a height of 1.36m). Thus, within group heterogeneity was apparent and was also consistent with literature reports of fascicle length heterogeneity (see Discussion). Additionally, we found a large range of fascicle lengths within each subject and muscle compartment. The average standard deviation of fascicle length within a single subject was 7.2mm for the anterior compartment, 6.7mm for the marginal compartment, and 6.3mm for the posterior compartment. While variability was evident within subjects, we did not find any significant differences in fascicle lengths between the CP and TD group.

PCSAs were significantly reduced among the CP cohort: mean deficits of 40.9% in PCSA (p = 0.014) (Fig 4). Normalized PCSA was reduced by 36.1 % in CP subjects (p = 0.001). A direct comparison between each age- and gender-matched subject’s PCSAs revealed a smaller PCSA than their TD counterpart.

**Fig 4:**
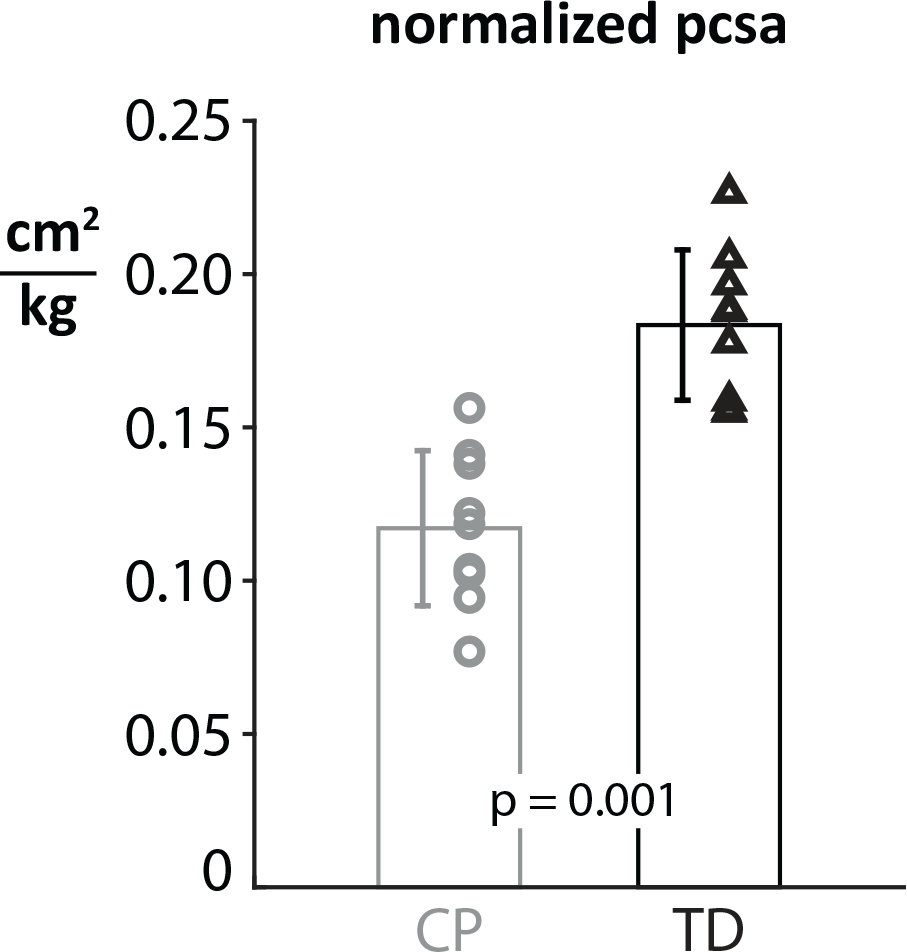
Normalized physiological cross-sectional areas (PCSAs) are significantly reduced in the CP cohort, indicating a reduced maximum force and functional weakness for this muscle. Here, PCSA was normalized by the body mass of each subject.

### Pennation Angle

The pennation angle is meant to describe how the muscle fibers are arranged and thus, how these arrangements influence the functionality. Yet, the computed angles for the selected characteristic fiber bundles varied widely between compartments and also within them (Fig 5). The angular range in the whole muscle ranged from 8.03° to 86.9°. Even within regions the ranges were large. As the pennation angle showed a lot of variations within and between compartments and also within and between the CP and TD group, no trend in reduced pennation angles was observed.

**Fig 5:**
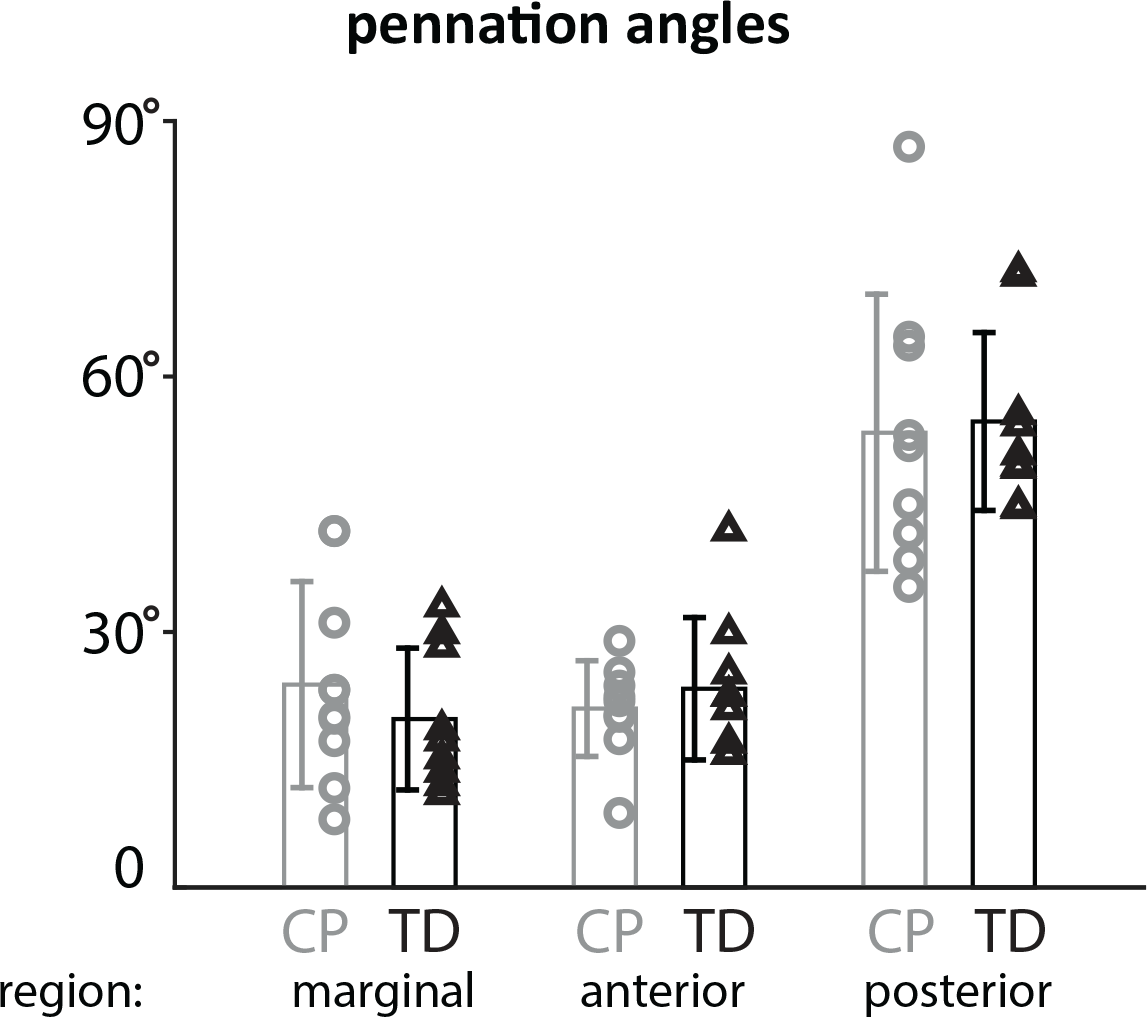
Pennation angles for all subjects in each compartment (marginal, posterior, anterior) show subject-specific results and large angle ranges also within the groups, but no significant differences between the groups.

## DISCUSSION

Deficits in muscle volume in CP have been previously observed and reported by many different groups(4,6,8,10,35,36). In the present study, we also observed significantly reduced muscle volumes. This was true for both absolute and body-size normalized muscle volumes. Deficits in muscle length and fascicle length in CP were not observed. Deficits in PCSA in this study were large in the CP group and consistent with literature findings for other lower limb muscles(11,37). Our reults suggest that volume deficits observed in CP are largely related to deficits in cross-section and indicate impaired strength capacity in CP.

From a geometrical perspective, muscle volume deficits may be related to deficits in fascicle length, PCSA, or both. Functionally, deficits in fascicle lengths are related to a reduced range of motion and contraction velocity while deficits in PCSA are related to reduced strength capacities. Based on the findings of this study, we conclude that volume deficits in CP are largely associated with decreased strength capacities rather than a reduced range of motion or reduced contraction velocities. Previously reported results on this question are somewhat varied: Handsfield et al.(6) did not determine fascicle lengths but reported reduced muscle lengths of 7.5% in the soleus in CP. Using ultrasound, Shortland et al.(38) and Barber et al.(11) found no fascicle length differences in the medial gastrocnemius between CP and TD children. Moreau et al.(39) found reduced fascicle lengths in the rectus femoris, but not the vastus lateralis, in CP. Differences between studies may reflect differences in the muscles analyzed and differences between cohorts or may result from methodological differences(17). The results found here, that fascicle lengths are not significantly different between CP and TD groups, have not been previously presented for the soleus muscle and is consistent with previous reports for other muscles in CP(11,14,38,40).

It should be noted that sarcomere lengths were unknown in this study. Sarcomere lengths represent a measure of length dependent force generation and passive stiffness and are used to calculate optimal fascicle length(14,41). Acquisition of sarcomere lengths *in vivo* requires the use of laser diffractometry or microendoscopy(42,43), which we did not have access to in this study. In a study using laser diffraction in adolescents with CP, Smith et al.(44) found sarcomere lengths to be about 17-22% longer in spastic hamstrings (gracilis and semitendinosus) muscles. A sarcomere length increase of 20% implies a 20% reduction in observed PCSA at the time of data acquisition. In our study we observed normalized PCSA deficits of 36%. In light of the potential differences in sarcomere lengths between CP and TD groups, it is possible that CP muscle fibers in our population may have fewer sarcomeres in series that are ‘stretched out’ to resemble the normal-length fibers in the TD population(14,44). Thus, although we did not examine sarcomere lengths in this study, when considered in the context of other studies on CP muscle architecture, it is possible that CP sarcomere lengths are longer than TD, optimal fascicle lengths are shorter than TD, and PCSAs are reduced compared to TD. The magnitudes of these effects suggest that PCSA deficit is greater than optimal fascicle length deficit. These results should be interpreted with caution, however, as the manifestation of CP is subject-specific and muscle-specific. Future studies that assess muscle volumes, fascicle lengths, and sarcomere lengths are warranted.

The definition of the PCSA used in this study(29,30) uses the muscle length to muscle fascicle length ratio, but not pennation angle. Another reported definition of PCSA(14) includes the pennation angle and muscle length to fascicle length ratio. While the former definition represents muscle cross-section perpendicular to muscle fiber direction, the latter definition is used to represent the cross-section scaled to a direction along the tendon axis. For our study, since the pennation angles were variable for each region of the soleus and also showed large heterogeneities within groups, it is difficult to define one representative pennation angle in order to compute the PCSA with respect to the pennation angle. This highlights a larger issue pertaining to defining 3D muscle architecture and comparing to 2D definitions from the literature. The soleus muscle has a complex organization comprising three aponeuroses and regions of muscle fibers with varying 3D orientations within and between regions. In the literature, muscles are often assigned a characteristic pennation angle to describe the orientation of fibers with respect to the tendon. Though useful for simply describing the angular offset between the tendon and muscle fibers, the 2D definition does not account for the complex organization of fibers in 3D space. Furthermore, when given the 3D vector descriptions of fiber directions within the muscle, it is difficult to decompose this information into a 2D angle. The right-hand side of Fig 6 illustrates a simplified representation of the principal fiber directions in sagittal view. Fibers are orientated in opposing directions, descending and ascending, on both sides of the anterior aponeurosis. Previous authors who used DTI have computed pennation angle as the minimum angle between an inserting fiber and the plane of the aponeurosis into which it inserts. This definition is also problematic as the minimum angle does not describe the azimuthal angular direction off of the aponeurosis in which the fiber is directed. In the case of a muscle with as complex an organization as the soleus, some consideration should be used in reporting and interpreting pennation angles. With regard to PCSA, the variety of pennation angles within the soleus muscle makes it difficult to compute one representative PCSA for the whole muscle where pennation angle is used as a multiplicative term. Thus, we used the definition of PCSA that omits consideration of pennation angle. More broadly, analysis of muscle function in terms of 3D orientation of fibers may benefit from tools which can account for the complex 3D orientations such as a finite element analysis(45,46).

**Fig 6:**
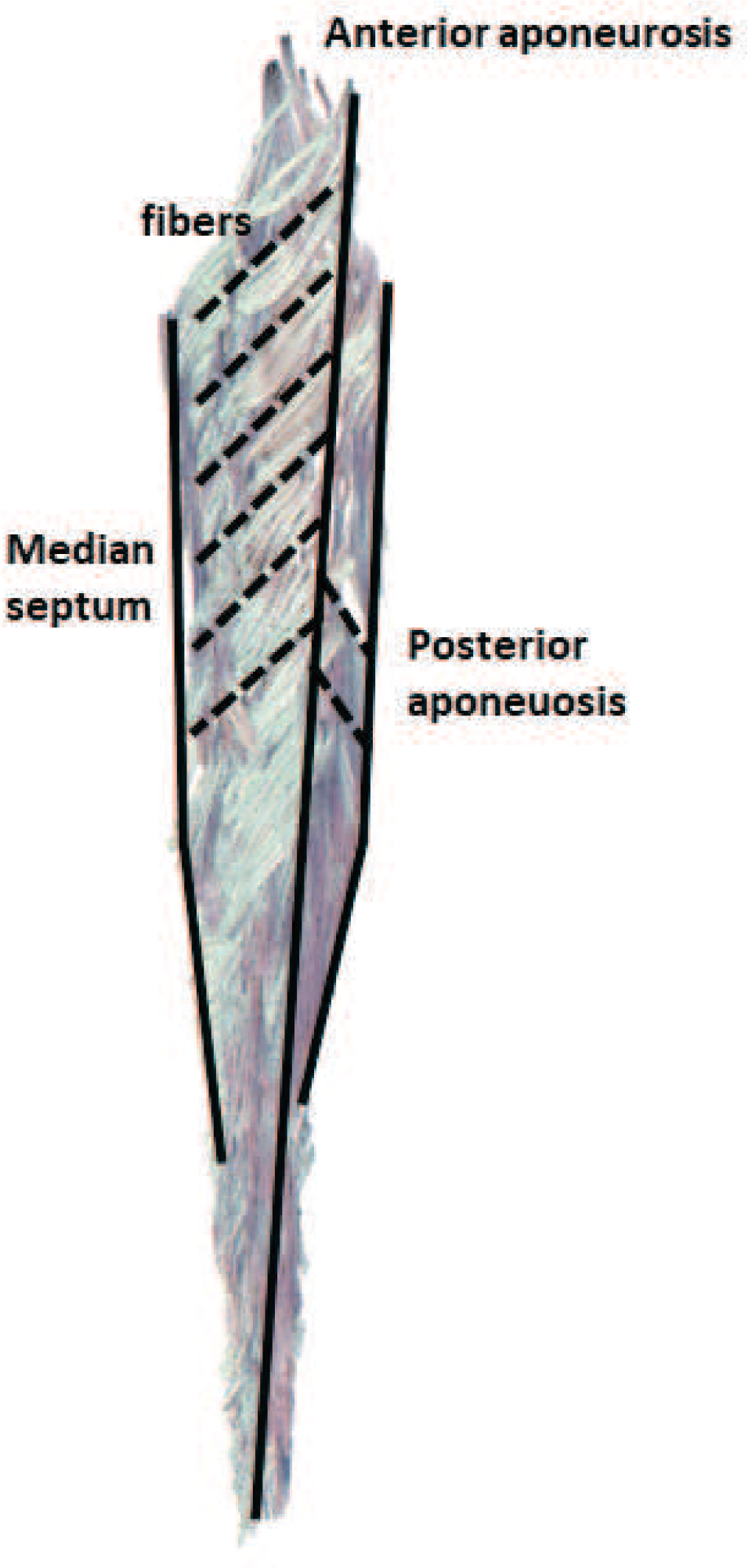
Complex soleus muscle architecture displayed from *in vivo* DTI data, mid-sagittal slice shown. The soleus muscle has three different aponeuroses where fibers originate and insert—the anterior aponeurosis, the median septum, and the posterior aponeurosis. Muscle fibers are oriented in opposite directions on either side of the anterior aponeurosis, creating an inverted “v” appearance.

The peak isometric force that a muscle is able to generate is directly related to the PCSA and the specific tension, σ, which is a measure of the force produced per unit area of skeletal muscle. This parameter has been estimated to be between 0.1 and 1.5 MPa(47,48) and is thought to vary somewhat between muscles and between individuals based on fiber type distribution and other factors. While specific tension may then be subject-specific, it is thought to be similar or reduced in subjects with CP compared to TD subjects. Taken together, a reduced PCSA in CP is representative of a direct reduction in peak isometric force. For the present study, isometric strength capacities in our CP cohort are expected to be reduced by at least the reduction in PCSA, but may be greater for subjects whose effective specific tension is also reduced.

There are several limitations to this study that bear consideration in addition to those discussed above. In this study, we imaged 9 individuals with cerebral palsy and 9 typically developing controls. This sample size is reasonable for an imaging study where data acquisition is expensive and time consuming; however, a larger population for this study would have conferred greater confidence in our results. Muscle length differences, particularly, showed a non-significant trend toward shorter fascicles in the cerebral palsy cohort. A greater number of participants may have revealed a significant difference here. As a technique, DTI does not specifically image muscle fibers—rather it is a technique for quantifying diffusion directions of water molecules in the tissue. By filtering the DTI data and implementing tractography algorithms that consider alignment of fibers and overall diffusion directions, muscle fiber tract directions can be reconstructed from DTI. While DTI and correct use of tractography algorithms has been shown to be robust and repeatable in determining muscle fiber directions(19–21,49), the nature of the method requires that users should not view each tract as a *de facto* muscle fiber. Rather the ensemble of tracts represents overall fiber directionality.

We divided the muscle into three compartments—posterior, anterior, and marginal—to be consistent with literature that investigated the soleus muscle(25,50). Considering the reconstructed fiber architectures using DTI, it seems possible that it bears defining other compartments as well. The large standard deviations even in small compartments of the soleus also implicate the diversity and complexity even within small spatial regions within the muscle. The large standard deviations we observed are consistent with literature reports where the muscle was divided into 32 compartments and carefully dissected(26). In these studies, the authors reported large standard deviations within compartments even despite the large number of compartments defined. These results notwithstanding, inspection of fiber directions from DTI in the soleus muscle for our set of subjects reveals that the major fiber orientations appear to be well-described by categorization into the anterior, posterior, and marginal compartments. Further analysis of or definition of alternative compartments may lead to a deeper understanding of this muscle and its function.

In this study, we used DTI and conventional MRI to determine muscle volumes, lengths, and fiber orientations in a CP and a TD population. We compared fascicle lengths and computed PCSA in the two groups, finding reduced volume and PCSA in CP. Though we did not compute sarcomere lengths in this study, the magnitude of PCSA deficits indicate that strength capacity is limited in CP and related to muscle architecture. Future studies may include a greater number of participants, utilize tools for determining sarcomere lengths, and assess additional lower limb muscles. Given the complex architecture of the soleus muscle, other tools may prove useful to understanding muscle architecture and its associated function in both cerebral palsy and typically developed populations. Mechanical simulations, for instance, may prove useful to understand how differences in muscle architecture and structure lead to altered function and mechanical behavior(45,46). An improved understanding of how muscles in CP function may motivate new therapies and help to predict their outcomes. The results of this study implicate muscle strengthening therapies as a potentially effective treatment regime. Physiotherapy, targeted strengthening, and habilitation therapies of muscles in CP may be very beneficial for children and adolescents with CP to strengthen their muscles, improve their muscle function, and improve their quality of life.

## Acknowledgements

This work was funded by The Wishbone Trust of the NZ Orthopaedic Association. The Whitaker Foundation and the Robertson Foundation funded personnel through fellowship support. We wish to acknowledge the contributions of Renee Miller, Beau Pontré, and the helpful staff of the Centre for Advanced MRI, especially Rachel Heron and Anna-Maria Lydon.

